# Privatization of biofilm matrix in structurally heterogeneous biofilms

**DOI:** 10.1101/742593

**Authors:** Simon B. Otto, Marivic Martin, Daniel Schäfer, Raimo Hartmann, Knut Drescher, Susanne Brix, Anna Dragoš, Ákos T. Kovács

**Affiliations:** Bacterial Interactions and Evolution Group, Department of Biotechnology and Biomedicine, Technical University of Denmark, 2800 Kongens Lyngby, Denmark; Terrestrial Biofilms Group, Institute of Microbiology, Friedrich Schiller University Jena, 07743 Jena, Germany; Max Planck Institute for Terrestrial Microbiology, 35043 Marburg, Germany; Department of Physics, Philipps-Universität Marburg, 35037 Marburg, Germany; Disease Systems Immunology Group, Department of Biotechnology and Biomedicine, Technical University of Denmark, 2800 Kongens Lyngby, Denmark

**Keywords:** *Bacillus subtilis*, biofilm, phenotypic heterogeneity, structural heterogeneity, exopolysaccharide

## Abstract

The self-produced biofilm provides beneficial protection for the enclosed cells, but the costly production of matrix components makes producer cells susceptible to cheating by non-producing individuals. Despite detrimental effects of non-producers, biofilms can be heterogeneous, with isogenic non-producers being a natural consequence of phenotypic differentiation processes. For instance, in *Bacillus subtilis* biofilm cells differ in the two major matrix components production, the amyloid fiber protein TasA and exopolysaccharides (EPS), demonstrating different expression levels of corresponding matrix genes. This raises questions regarding matrix gene expression dynamics during biofilm development and the impact of phenotypic non-producers on biofilm robustness. Here, we show that biofilms are structurally heterogeneous and can be separated into strongly and weakly associated clusters. We reveal that spatiotemporal changes in structural heterogeneity correlate with matrix gene expression, with TasA playing a key role in biofilm integrity and timing of development. We show that the matrix remains partially privatized by the producer subpopulation, where cells tightly stick together even when exposed to shear stress. Our results support previous findings on the existence of ‘weak points’ in seemingly robust biofilms as well as on the key role of linkage proteins in biofilm formation. Furthermore, we provide a starting point for investigating the privatization of common goods within isogenic populations.

**IMPORTANCE:** Biofilms are communities of bacteria protected by a self-produced extracellular matrix. The detrimental effects of non-producing individuals on biofilm development raises questions about the dynamics between community members, especially when isogenic non-producers exist within wild-type populations. We asked ourselves whether phenotypic non-producers impact biofilm robustness, and where and when this heterogeneity of matrix gene expression occurs. Based on our results we propose that the matrix remains partly privatized by the producing subpopulation, since producing cells stick together when exposed to shear stress. The important role of linkage proteins in robustness and development of the structurally heterogeneous biofilm provides an entry into studying the privatization of common goods within isogenic populations.

## INTRODUCTION

Biofilms are communities of tightly associated microorganisms encased in a self-produced extracellular matrix (1). This matrix provides shielding against biotic factors, such as antibiotics (2, 3), natural competitors or predators (4, 5) and abiotic factors, such as harsh physicochemical (6) or shear stress (7). As components of the biofilm matrix are costly to produce and they can be shared within the population, matrix producers are potentially susceptible to social cheating, where non-producing mutants benefit from productive community members (8–10). This ‘tragedy of the commons’ principle, in which non-participating users cannot be excluded from the use of common goods (9, 11, 12), has already been demonstrated for *Pseudomonas fluorescens* SBW25, for which exploitation by an evolved non-producer resulted in biofilm collapse (13). Alternatively, production of the matrix components may not be easily exploitable if there is limited sharing, low cost of production, or spatial assortment of cells within the biofilm (14, 15). Finally, long-term cheating on matrix production may have evolutionary consequences for the producers, changing the phenotypic heterogeneity pattern of matrix expression within the population (16).

Although so-called ‘cheating’ is traditionally associated with loss-of-function mutation in matrix genes, phenotypic non-producers (cells in the so-called OFF state) can be an intrinsic part of clonal wild-type populations (17–19). For instance in *Bacillus subtilis* NCBI 3610, a member of a probiotic and plant growth-promoting species (20, 21), the aforementioned phenotypic heterogeneity is fundamental to biofilm development, with individual cells exhibiting different tendencies to differentiate or express motility determinants (22, 23). Formation of pellicle biofilms, also referred to as ‘liquid-air interface’ biofilms, in *B. subtilis* includes aerotaxis-driven swimming towards the liquid-air interface, subsequent motility loss and adherent extracellular matrix production (24, 25). This differentiation of motile cells, becoming matrix-producing cells and spores, is not terminal, with genetically identical cells being able to alter their gene expression (26). While exploitability of the extracellular matrix by non-producing mutants has been extensively studied, social interactions between clonal matrix producers and non-producers in biofilms have been explored less. According to Hamilton’s rule, altruistic sharing of public goods can easily evolve within isogenic populations, by means of inclusive fitness benefits (8). In other words, as long as the recipient carries the cooperative gene, cooperation should be evolutionary stable in the absence of additional stabilizing mechanisms (27–30).

Still, it is not clear to what extent the matrix is shared between phenotypically heterogeneous producers and non-producers, whether the presence of a non-producing subpopulation has consequences for local biofilm robustness and if/how the distribution of non-producers changes during biofilm development. In fact, biofilms are non-uniform structures with variable local cell and polymer densities (31), which could be linked to different behavior of cells within a clonal population. Understanding how the heterogeneity of gene expression is linked to both biofilm development and structural robustness would provide better insight into the dynamics of biofilm communities.

The extracellular matrix of *B. subtilis* NCBI 3610 consists of two major components: an amyloid protein, TasA, and exopolysaccharide (EPS) (32). The EPS component is synthesized by protein products encoded by the *epsA-epsO* operon, with Δ*eps* mutants producing a weak and fragile biofilm (32, 33). The protein component TasA forms amyloid fiber-like structures (34, 35) and it is encoded in the *tapA*-*sipW*-*tasA* operon, with Δ*tasA* mutants unable to produce a biofilm (36). Mutant strains of *B. subtilis* NCBI 3610 lacking both operons cannot form a biofilm, whereas strains producing one of the components can complement each other and produce a wild-type-like pellicle (15, 32, 37). In this study, we demonstrate that, under exposure to shear stress, these seemingly sturdy pellicle biofilms disintegrate into extremely robust aggregates and single cells. We reveal that spatial and temporal changes in biofilm structural heterogeneity correlate with changes in expression of biofilm components, as cells in the ON state dominate within unbreakable biofilm aggregates. Therefore, despite inclusive fitness benefits from sharing the public goods within an isogenic population of producers, phenotypic cooperators (ON cells) still partially privatize the biofilm matrix. Further, we propose that the protein matrix component TasA plays a key role in maintaining biofilm robustness, with major consequences for the timing of development and the overall productivity of the biofilm. In general, our study links changes of phenotypic heterogeneity pattern with different stages of biofilm formation and reveals a fingerprint of such heterogeneity in biofilm structural robustness. It also reveals that privatization of public goods occurs even in isogenic microbial populations.

## RESULTS

### Structural heterogeneity develops in late stages of pellicle growth

We began from a simple question: does phenotypic heterogeneity of matrix gene expression in *B. subtilis* (16, 19, 37) translate into non-uniform robustness within the biofilm? We will refer to such non-uniform biofilm robustness as “structural heterogeneity”. Biofilms were mechanically disrupted by vortexing with sterile glass sand (Fig. 1a). Consequently, biofilm cells could be separated into two fractions: a fragile dispersible fraction, and a robust non-dispersible fraction, of which ‘clumps’ could be easily observed under a microscope with low magnification and persisted for up to 8 days old pellicles. (Fig. 1b). This structural heterogeneity of the biofilm was predominantly observed in mature pellicles (older than 24h) and the fractions were dynamic as the pellicle aged (Fig. 1c). During the establishment of a pellicle, around 13–16 h after inoculation of the bacteria into MSgg medium, roughly 0.25-0.32 fraction of cells could be assigned to fragile fraction. Between 19 and 24 h, the juvenile pellicle was mostly structurally homogenous, as it consist solely of a robust fraction. In later stages of pellicle development, the biofilm was again structurally heterogeneous, with an increase in cell number in the fragile fraction (Fig. 1c). To better understand the interplay between the robust and fragile fractions of biofilm during its development, we looked into changes in absolute cell numbers in both fractions (Fig. 1d). At most stages of biofilm development the amount of cells in robust and fragile fractions differed significantly (Fig. 1d). Moreover, biofilm development coupled with an increase of biofilm biomass could be divided into 2 stages: 1-early biofilm development (between 16-21 hours) which characterized with a dramatic increase in total number of cells in robust fraction (and only moderate increase of cells in fragile fraction); 2-late biofilm development (between 24-48 hours) when the number of cells in robust fraction remain constant and cells in the fragile fraction largely increased in numbers (Fig. 1d). Overall, these results indicate that structural heterogeneity in mature pellicles (older than 24h) results from the emergence of loosely attached (fragile) population of cells on the ‘backbone’ of robust cells. Turning point between early and late biofilm development takes place after around 24 hours.

**FIG 1.**
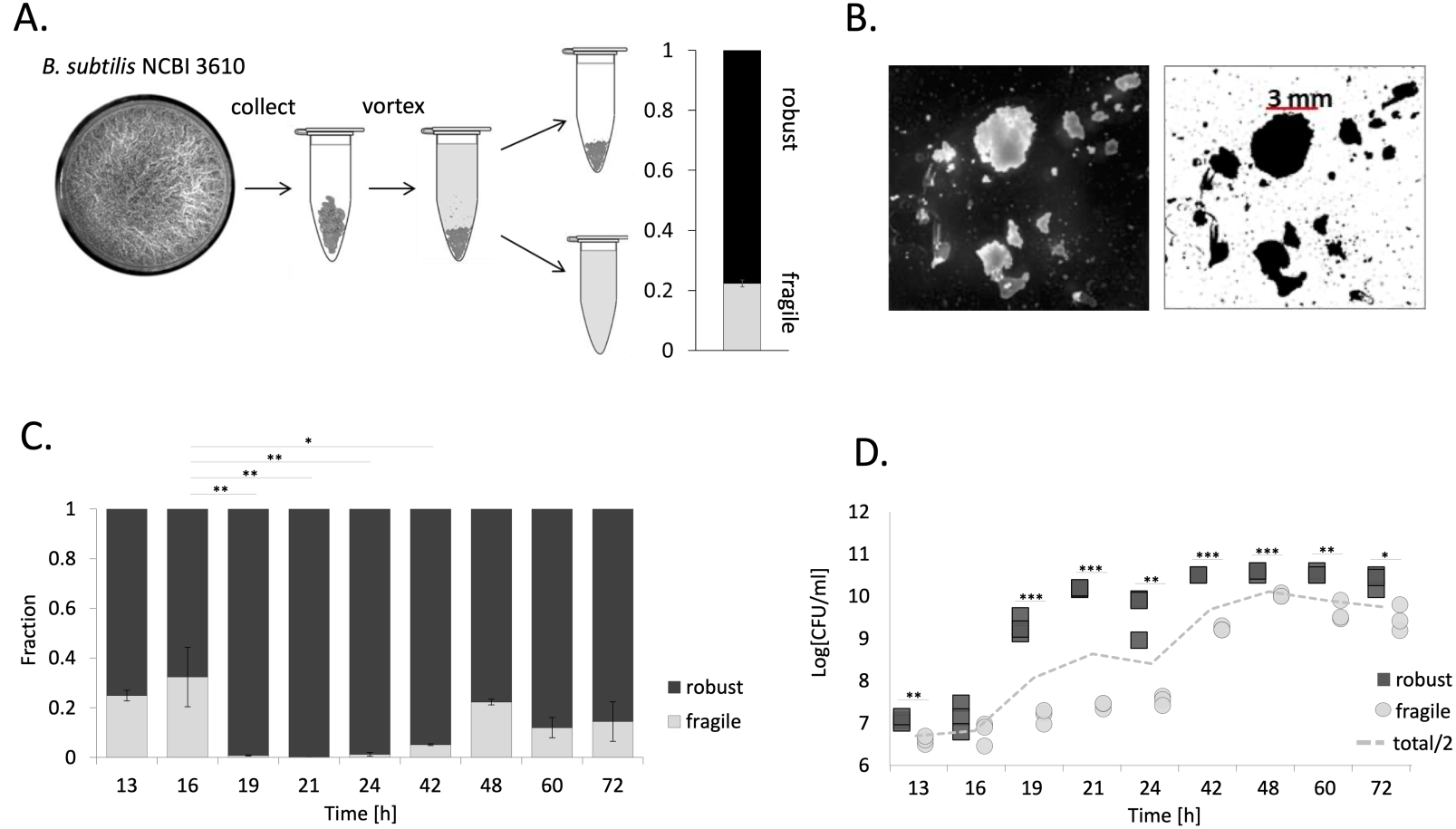
Structural heterogeneity in pellicle biofilms. (a) Mechanical disruption of biofilms into “robust” and “fragile” fractions by vortexing the pellicle with sterile glass beads. (b) Microscopy images show that the robust fraction consists of non-dispersible clumps that could be observed under a microscope with low magnification. These clumps were also present in 8-day-old pellicles. Scale bars indicate 5 mm. (c) Temporal changes in relative abundance of cells belonging to the robust and fragile fraction of the biofilm. The dark grey bar represents the robust fraction, while the light grey bar represents the fragile fraction. Data represent an average from biological triplicates and error bars correspond to standard errors. (d) Changes in total cell counts in biofilm (dashed grey line) and cells in robust and fragile fraction over time were represented in logarithmic units. All data points were shown on the graph. For panels c and d, * stands for p<0.05, ** for p<0.01, *** for p<0.001.

### Temporal changes in structural heterogeneity overlap with changes in phenotypic heterogeneity

Having shown that structural heterogeneity changes throughout biofilm development, we next sought to determine what underlies these changes. Considering that biofilms are non-uniform structures with variable polymer densities, we chose to investigate the expression of the *epsA-epsO* and *tapA-sipW-tasA* operons encoding the two major components of the biofilm, EPS and amyloid protein TasA, respectively (31, 32). Transcription levels were analyzed by flow cytometry using P_*eps*_*-gfp* and P_*tapA*_*-gfp* reporter strains at various pellicle ages (Fig. 2). Expression of both P_*eps*_*-gfp* and P_*tapA*_*-gfp* was shown to be low at 12 h (in most replicates insignificantly different from control, non-labelled strain), when pellicles first emerged, indicating that most cells produced no or very low amounts of EPS and TasA. In both strains (P_*eps*_*-gfp* and P_*tapA*_*-gfp*), one replicate showed an emergence of an ON-subpopulation, indicating biological stochasticity at this very early timepoint (Fig. 2, Dataset S11). The relative size of the ON-subpopulation increased significantly between 12 and 16h in both P_*eps*_*-gfp* and P_*tapA*_*-gfp*, thus in the majority of cells; 61% in P_*eps*_*-gfp*, and 66% in P_*tapA*_*-gfp* (Fig. 2, Dataset S1). Further changes were observed between 16 and 20h, as the mean *eps* expression intensity increased significantly, so that OFF-subpopulation became the low-*eps* subpopulation (fluorescent signal significantly increased above the control level), and ON-subpopulation shifted towards a higher expression level. At the same time, the expression pattern of *tasA* became unimodal – with the OFF-subpopulation disappearing completely (Fig. 2, Dataset S1). Between 20-24h, the *eps*-expressing subpopulation further increased in size, while opposite was observed for the *tasA*-expression pattern, where the OFF-subpopulation reappeared again (Fig. 2, Dataset S1). At later time points, the heterogeneity level in both P_*eps*_*-gfp* and P_*tapA*_*-gfp* increased once again, with more pronounced OFF-subpopulations. In mature pellicles (older than 24h), similarly to the onset of biofilm formation, OFF subpopulations were in the majority (Fig. 2, Dataset S1).

**FIG 2.**
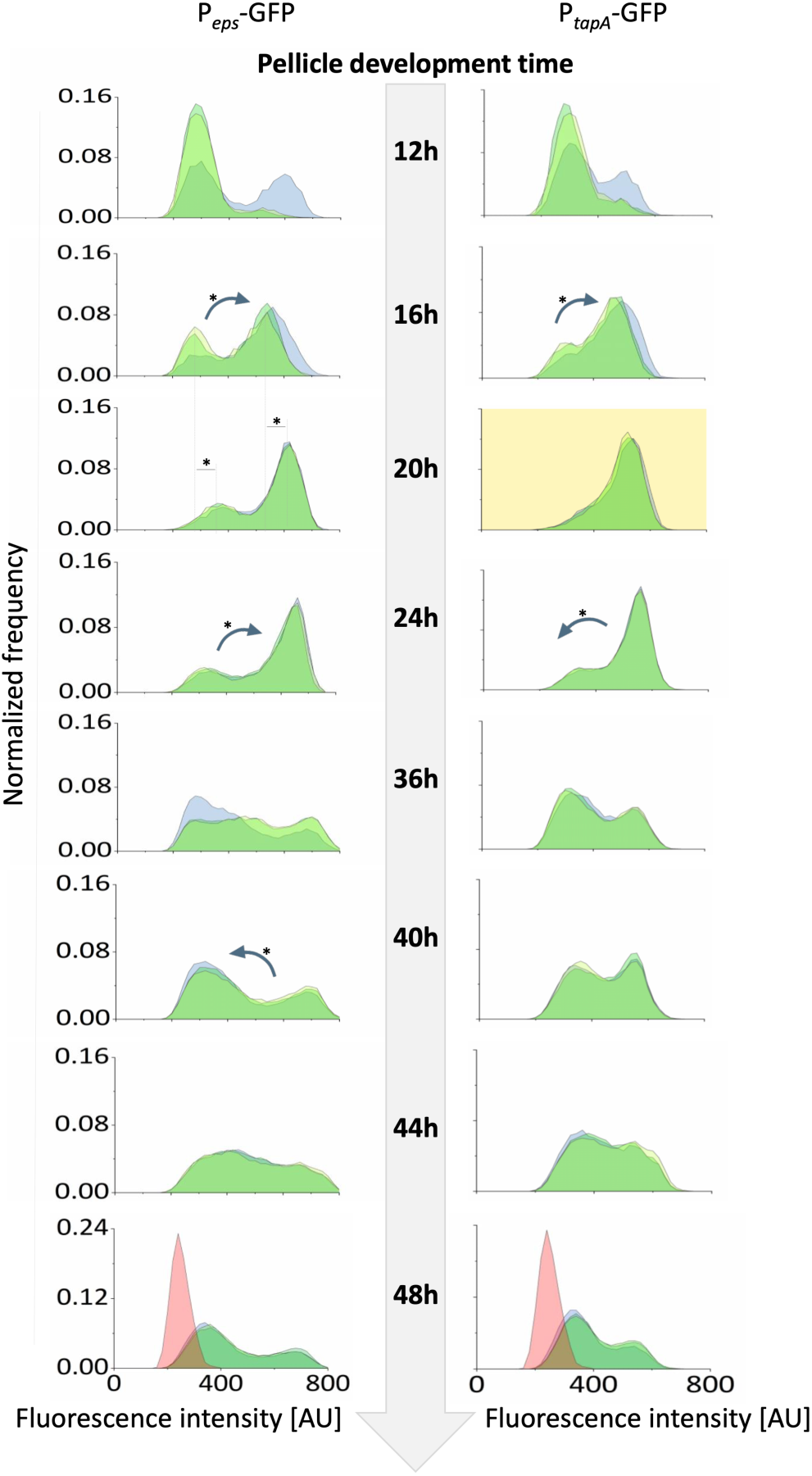
Changes in matrix gene expression during biofilm development assessed by Flow cytometry. Flow cytometry analysis showing distributions of fluorescence intensities of GFP-based transcriptional reporters for *epsA-epsO* (left) and *tapA-sipW-tasA* (right) at various time points throughout biofilm development. Histograms obtained for all biological replicates (n=3) are overlayed for each timepoint. Data where distribution of matrix gene expression was unimodal (P_*tapA*_-GFP, 20h) is marked with yellow background. Significant shifts of mean expression level in each subpopulations were indicated by dashed lines and asterisk. Significant changes in relative size of subpopulation with low- and high-matrix gene expression, were shown as arrows (pointing towards shift direction) and asterisk. For changes in mean expression and subpopulation relative size, only significant differences between 2 neighboring timepoints were depicted on the image. Data for non-labelled control were acquired for 48h-old pellicle and integrated into the corresponding histograms as a red overlay.

We observed similar changes in phenotypic heterogeneity when P_*eps*_*-gfp* strains were analyzed under a confocal microscope (Fig. S1). Expression of *epsA-epsO* was most prevalent from 19-24 h with OFF subpopulations being observed at earlier and later time points. As images derived from intact biofilms, that contain clusters of ON and OFF cells, the bimodality of *eps* expression manifested after overlay of data from several frames per timepoint (Fig. S1). Overall, changes observed in phenotypic heterogeneity of *epsA-epsO* and *tapA-sipW-tasA* expression correlated with the temporal changes we observed in structural heterogeneity of the biofilm. The so called ‘turning point’ in biofilm development, where growth of robust fraction stops and growth of fragile fraction begins (Fig. 1d) overlaps with a switch of *tasA* expression from unimodal ON state to bimodality, and increasing numbers of *eps*-expressing cells. Late stage of biofilm development, when fragile fraction increases in numbers (Fig. 1d), overlaps with an increase in relative sizes of OFF-subpopulations with respect to both *eps* and *tasA*. These coupling between temporal changes in biofilm structural heterogeneity and matrix genes expression led us to hypothesize a spatial correlation between ON cells and the non-dispersible parts of the biofilm.

### Expression of *epsA-epsO* and *tapA-sipW-tasA* operons in robust and fragile fractions

To investigate if robustness is spatially related to high levels of polysaccharide and amyloid protein production, pellicles established by P_*eps*_*-gfp* and P_*tapA*_*-gfp* reporter strains were mechanically disrupted after which ‘clumps’ and dispersible fractions were separately analyzed by flow cytometry (Fig. 3, Dataset S1). Pellicle of ages of 24, 36 and 48 h were chosen because of the shift towards phenotypic heterogeneity we had observed as the pellicles aged (Fig. 2, Fig. S1). We noted that already at 24 h, there was a slight difference in *epsA-O* expression pattern between the robust and fragile pellicle fraction, as the percentage of ON-cells was significantly larger in the robust fraction (Fig. 3a, Dataset S1). After 36 hours, the robust fraction of P_*eps*_*-gfp* strain not only contained higher percentage of ON-cells compared to the fragile fraction, but also the *epsA-O* expression levels in OFF subpopulation increased beyond the background noise, shifting the ON/OFF distribution towards a low ON/high ON scenario (Fig. 3a, Dataset S1). After 48 hours, major changes took place in the robust fraction of P_*eps*_*-gfp*, where relative number of high ON cells decreased and low ON subpopulation shifted back to an OFF state (Fig. 3a, Dataset S1). In contrast to P_*eps*_*-gfp*, major differences between the robust and fragile fractions of P_*tapA*_*-gfp* were observed in late biofilms (after 48 hours), where the robust fraction of the biofilm still contained substantial amount of ON cells, with significantly higher expression levels compared to those observed in the fragile fraction of the biofilm (Fig. 3b, Dataset S1).

**FIG 3.**
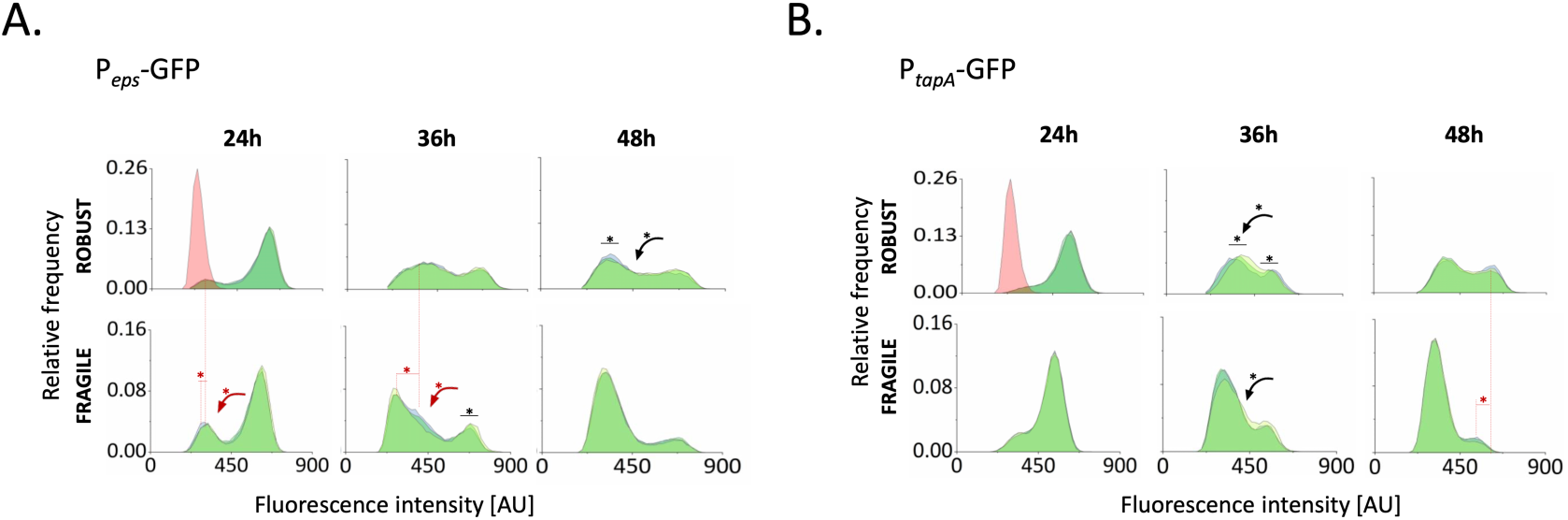
Expression of matrix genes in robust and fragile fractions of the biofilm. (a) Confocal microscopy images of intact and mechanically disrupted P_*eps*_*-gfp*, P_hyperspank_-*mKATE2* pellicles. Images show the overlay of magenta and green channels. Scale bar indicates 35 µm. (b) Flow cytometry analysis showing average (*n* = 3) distributions of fluorescence intensities of mechanically disrupted P_*eps*_*-gfp* and P_*tapA*_*-gfp* reporter strains after 24, 36, and 48 h. The blue histogram represents the robust fraction, while yellow graph represents the fragile fraction; grey graph depicts non-labelled cells. Data for non-labelled control were acquired for 48h-old pellicle and integrated into the upper left histograms of left and right panels, as a red overlay.

Overall, this analysis suggests that in early biofilm, fragile and robust fraction differ mostly in *eps* expression pattern (Fig. 3a, Dataset S1). On the other hand, in mature biofilms, when structural heterogeneity becomes more pronounced due to the increasing size of the fragile fraction (Fig. 1c, d), the robustness seems to be maintained through high levels of *tasA* expression (Fig. 3b, Dataset S1).

Additionally, we observed TasA non-producers, cocultured with EPS non-producers, to be dominant at the breakage points of clumps (Fig. S2), suggesting an involvement of TasA in biofilm integrity. Although our preliminary observation of increased abundance of Δ*tasA* mutant at the pellicle breakage points requires further studies, it further points towards importance of the TasA protein in biofilm mechanical robustness.

### TasA non-producers have negative effects on the timing of pellicle development and final pellicle productivity

Next, we aimed to determine how each matrix component affects biofilm development. First, we competed biofilm mutants lacking one or both matrix components against the wild-type in competition assays with 1:1 relative inoculation. Relative fitness of biofilm mutants in both the liquid medium was assessed after 24 and 48 h. Although Δ*eps* mutant and wild-type strains were equally fit in the pellicle, the mutant could outcompete wild-type in the liquid medium (below the pellicle biofilm) (Fig. 4a). On the contrary, the Δ*tasA* mutant was clearly losing the competition against the wild-type in the pellicle (Fig. 4a). Furthermore, the Δ*eps*Δ*tasA* mutant was significantly outcompeted in the pellicle and in the liquid after 48 h (Fig. 4a) which was clearly evident from the microscopy images of mixed pellicles (Fig. S3)

**FIG 4.**
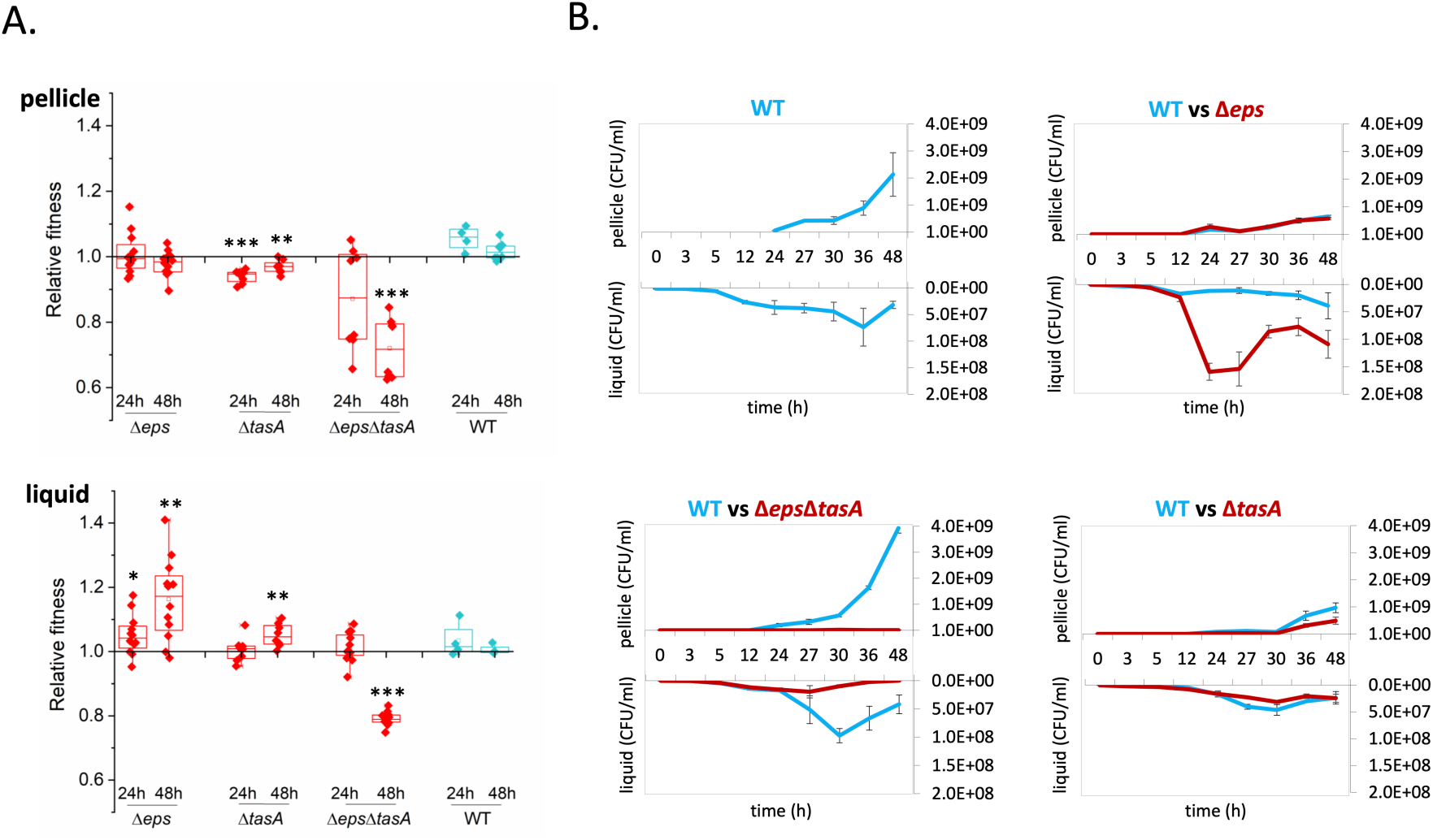
Fitness of biofilm mutants in the pellicle and their effect on biofilm development. (a) Relative fitness of biofilm mutants in the pellicle biofilm (robust + fragile fraction) and in liquid medium (below the biofilm) measured after 24 h and 48 h based on total colony-forming units (CFU)/ml counts. Boxes represent Q1–Q3, lines represent the median, and bars span from max to min. *indicates p<0.05; **p<0.01; ***p<0.001. (b) Temporal changes in productivity during biofilm development in wild-type monoculture, and in cocultures of the wild-type with either Δ*eps*, Δ*tasA* or Δ*eps*Δ*tasA* strains. Productivity was assessed at different time points both in the pellicle and in liquid medium (below the biofilm). Pellicles were collected, resuspended in 1ml of saline solution, disrupted and serially diluted for CFU assay – CFU/ml stands for the number of cells obtained after pellicle disruption/1ml of saline solution. Data points represents the average of *n* = 3 biological replicates and error bars correspond to standard error.

The reduced performance of the Δ*tasA* mutants in the pellicle suggests it has negative effects on biofilm development. Thus, the effect of biofilm mutants on pellicle productivity (i.e. total number of cells in the pellicle) during development was assessed (Fig. 4b, Dataset S1). Cocultures of wild-type and biofilm mutants were mixed 1:1, and colony-forming unit (CFU) productivity in the liquid and pellicle was determined at various time points throughout the development. We noted that both Δ*eps* and *ΔtasA* significantly slowed down pellicle development, which was not the case for Δ*eps*Δ*tasA* (Fig. 4b, Dataset S1). The effect was especially pronounced for *ΔtasA* (Fig. 4b) and could also be captured by stereomicroscope time lapse movies (Fig. S4 and Video S1). In conclusion, EPS, and especially TasA non-producers seem to slow down pellicle development and reduce final pellicle productivity (Fig. 4b, Fig. S4 and Video S1). In addition, lack of a negative impact of Δ*eps*Δ*tasA* suggests that at least one of the two matrix components is required for positioning of the biofilm mutant in the pellicle and its negative effects on development and productivity.

### TasA non-producers diminish pellicle robustness, while EPS non-producers do not

The function of TasA as a linkage protein and the importance of TasA for pellicle development suggests its significant contribution towards pellicle robustness. To investigate this, cocultures containing increasing percentages of Δ*eps* or Δ*tasA* were mixed with the wild-type and CFU productivity in the robust and fragile fractions of the pellicle was determined (Fig. 5). When the wild-type was confounded with Δ*eps*, wild-type productivity in the robust fraction was reduced but its level was maintained as the proportion of Δ*eps* increased. Consistently, in both fragile and robust fraction we detected significant negative correlation between amounts of the WT and Δ*eps* (Pearson corr. = -0.85, p<1.2×10^-6^; -0.61, p<0.004; for fragile and robust fraction, respectively) suggesting that in both fractions, the mutant was able to compete with the WT (Fig. 5). Nevertheless, we did not detect significant correlation between ratio of Δ*eps* and biofilm robustness (Pearson corr. = 0.11, p<0.61). In contrast, if the wild-type was mixed with Δ*tasA*, it failed to incorporate into the robust pellicle fraction (Pearson corr. = 0.16, p<0.46), however it turned out to be detrimental for biofilm robustness (Pearson corr. = -0.85, p<1.6×10^-6^). These results clearly show the negative impact of TasA non-producers on pellicle robustness, and the importance of TasA for incorporation into robust part of the biofilm.

**FIG 5.**
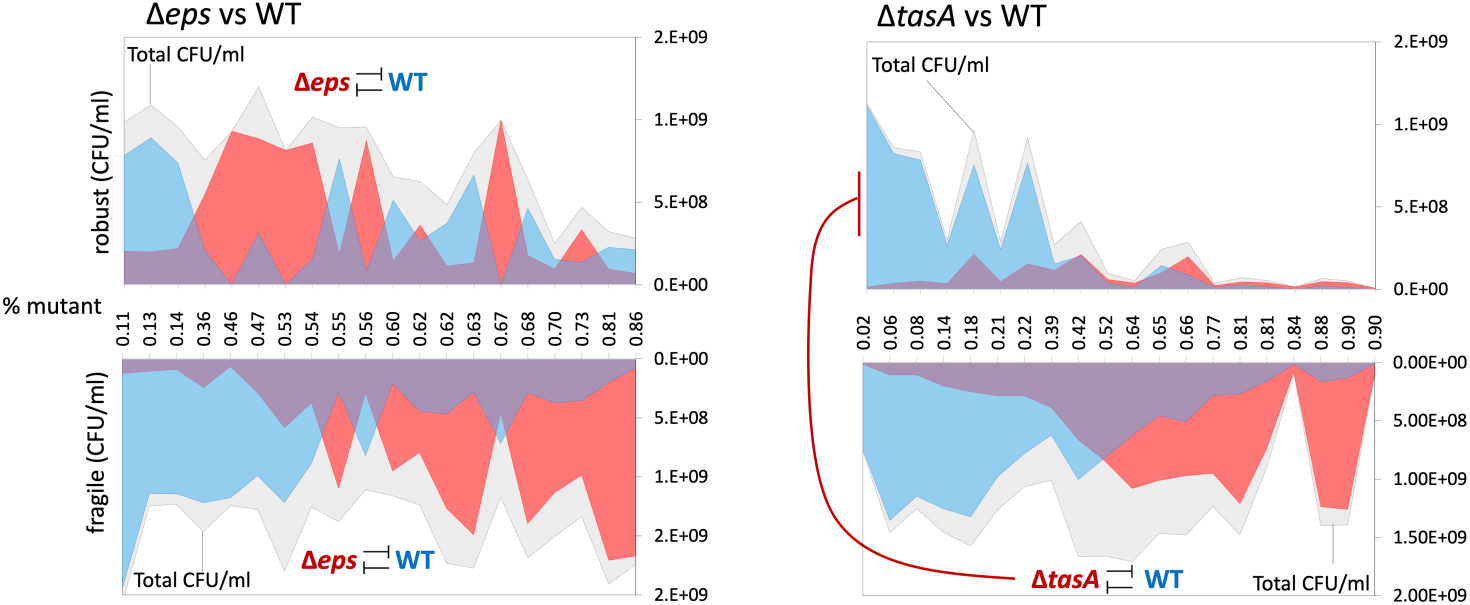
Effect of biofilm mutants on pellicle robustness. Productivities of wild-type and mutants based on total CFU/ml were assessed in mechanically disrupted robust and fragile fractions in cocultures of WT with increasing ratios of either Δ*eps* or Δ*tasA*. Relationships between WT and mutants were examined using Pearson correlation coefficient. Significant negative correlations between WT and mutants, or between mutant and size of robust fraction, are labelled by inhibition symbol.

## DISCUSSION

Studies of the social interactions between genetically engineered matrix producers and non-producers have become a common approach in sociobiology of biofilm communities (14, 37–39). Here, we addressed the consequences of native within-population phenotypic heterogeneity in matrix production for robustness, productivity and timing of biofilm development. We revealed that production of matrix components shifts throughout biofilm development and that these changes correlate with temporal and spatial changes in biofilm robustness.

Biofilm development can be studied from different aspects (40–43). Here, we showed that in the initial stage of biofilm formation, the majority of the population was in an ON state, followed by heterogeneity in older biofilms. We show that in the timeframe between 16-20 hours, where an increase of robust pellicle biomass is the most pronounced (Fig 1D), there is a significant shift *eps*-expression intensity and switching ON of *tasA* expression in nearly all biofilm cells. Therefore, our results link temporal dynamics in matrix genes expression with temporal changes of robust biofilm biomass.

This data in line with previous studies, in which the spatiotemporal dynamics patterns of gene expression during the formation of submerged *Escherichia coli* biofilms were investigated (40). Moreover, bimodal expression of curli fibers was demonstrated, with high curli expression being confined to dense cell aggregates. The bimodal spatial expression of the structurally important curli fibers suggests a similar role for TasA, with these higher cell density aggregates providing protection against shear stress. Furthermore, in another study, the production of curli fibers was shown to protect the biofilm population against bacteriophage (41).

Importantly, cells exhibiting the ON state are more likely to occupy the most robust areas of the biofilm, thereby privatizing the benefit from matrix production under exposure to shear stress. Previous studies have shown that phenotypic heterogeneity of matrix production is present in biofilms of different species (17–19), with a similar phenomenon likely to occur in other biofilm-producing bacteria. Recently, quantitative visualization of *Pseudomonas aeruginosa* PAO1 aggregates has shown peak alginate gene expression in cells proximal to the surface compared with cells in the interior (44). Although it is likely that the interior of the *B. subtilis* pellicle biofilm contains more OFF cells, we believe that the temporal shift we observed from heterogeneity to homogeneity, then again towards heterogeneity is due to phenotypic differences between randomly distributed isogenic cells.

Accordingly, we revealed that TasA non-producers have adverse effects on the timing of matrix development, productivity and robustness, which was not the case for the EPS non-producers. As EPS is likely costlier to produce and less privatized than TasA (37), the diminishing effects of Δ*tasA* may be linked to the specific structural role of TasA in the matrix, as could also be supported by its distinct localization pattern (45). The dominance of ON cells in the robust biofilm fraction was especially pronounced for *tasA* expression. Furthermore, we observed TasA-non-producers to be dominant at breakage points of biofilm clumps, suggesting these areas are weak points in biofilm integrity.

Conceivably, TasA functions similarly to the structural protein RmbA described in *Vibrio cholerae* biofilms, creating strong linkage between the producing cells (39). If the linkage role holds true, Δ*tasA* cells should be impaired in their ability to integrate into pre-established wild-type pellicles, which will be explored in the future. TasA was shown to have a strong adhesive role during interspecies interactions (46) and has been linked to structural integrity and physiology of *B. subtilis* biofilms (47, 48).

Our results support previous observations showing the importance of linkage proteins in formation of biofilms (32), as well as the presence of non-uniform biofilm structures (31). It remains to be discovered how the extracellular matrix remains privatized by ON cells and what are the ecological consequences or potential evolutionary benefits from biofilm structural heterogeneity. One possibility is bet-hedging where weakly associated cells would adapt for short starvation periods, while early-sporulating aggregates are adapted for longer starvation periods, as proposed for slime molds (49). It remains to be tested, whether robust and fragile fractions of *B. subtilis* biofilms differ in sporulation dynamics.

Our work has four major conclusions: 1) seemingly integral biofilms consist of robust and loosely associated cells, thereby being structurally heterogeneous; 2) changes in the phenotypic heterogeneity pattern of matrix gene expression correlate with changes in biofilm structural heterogeneity in time and space; 3) TasA non-producers have detrimental effects on matrix development and structural integrity; 4) even in clonal microbial populations, where cooperation is stabilized by inclusive fitness benefits, public goods may be partially privatized by phenotypic producers.

## MATERIALS AND METHODS

### Bacterial strains and cultivation methods

Strains used in this study are listed in Table 1. All strains were maintained in lysogeny broth (LB; LB-Lennox, Carl Roth), while MSgg medium (5 mM potassium phosphate buffer (pH 7), 100 mM MOPS (pH 7), 2 mM MgCl_2_, 700 μM CaCl_2_, 50 μM MnCl_2_, 50 μM FeCl_3_, 1 μM ZnCl_2_, 2 μM thiamine, 0.5% (v/v) glycerol, 0.5% (w/v) glutamate) was used to induce biofilm formation (24). To obtain pellicle biofilms, bacteria were grown in static liquid MSgg medium at 30°C for 48 h, using 1% inoculum from overnight cultures. Prior to experiments, pellicles were sonicated according to an optimized protocol that allows for disruption of biofilms without affecting cell viability (15, 50). Productivity was determined by plating dilutions on LB-agar to obtain CFU.

**Table 1.** Strains used in this study

| Strain | Genotype | Reference |
| --- | --- | --- |
| DK1042 | 3610 *comI*<sup>Q12I</sup> (wild type) | (55) |
| TB34 | DK1042 but *amyE*::P<sub>hyperspank</sub>-*gfp* (Cm<sup>R</sup>) | (56) |
| TB35 | DK1042 but *amyE*::P<sub>hyperspank</sub>-*mKate2* (Cm<sup>R</sup>) | (57) |
| TB500 | DK1042 but *amyE*::P<sub>hyperspank</sub>-*gfp* (Spec<sup>R</sup>) | (56) |
| TB501 | DK1042 but *amyE*::P<sub>hyperspank</sub>-*mKate2* (Spec<sup>R</sup>) | (58) |
| TB514 | DK1042 but *eps*::Tet<sup>R</sup>, *tasA*::Spec<sup>R</sup>, *amyE*::P<sub>hyperspank</sub>-*gfp* (Cm<sup>R</sup>) | This study |
| TB515 | DK1042 but *eps*::Tet<sup>R</sup>, *tasA*::Spec<sup>R</sup>, *amyE*::P<sub>hyperspank</sub>-*mKate2* (Cm<sup>R</sup>) | This study |
| TB524 | DK1042 but *eps*::Tet<sup>R</sup>, *amyE*::P<sub>hyperspank</sub>-*gfp* (Spec<sup>R</sup>) | (58) |
| TB525 | DK1042 but *eps*::Tet<sup>R</sup>, *amyE*::P<sub>hyperspank</sub>-*mKate2* (Spec<sup>R</sup>) | (58) |
| TB538 | DK1042 but *tasA*::Km<sup>R</sup>, *amyE*::P<sub>hyperspank</sub>-*gfp* (Spec<sup>R</sup>) | (58) |
| TB539 | DK1042 but *tasA*::Km<sup>R</sup>, *amyE*::P<sub>hyperspank</sub>-*mKate2* (Spec<sup>R</sup>) | (58) |
| TB601 | DK1042 but *eps*::Tet<sup>R</sup> | (50) |
| TB602 | DK1042 but *tasA*::Spec<sup>R</sup> | (56) |
| TB852 | DK1042 but *eps*::Tet<sup>R</sup>, *tasA*::Km<sup>R</sup> | This study |
| TB863 | DK1042 but *tasA*::Km<sup>R</sup> | (37) |
| TB864 | DK1042 but *amyE*::P<sub>hyperspank</sub>-*mKate2* (Cm<sup>R</sup>) *sacA*::P<sub>eps</sub>-*gfp* (Km<sup>R</sup>) | (50) |
| TB865 | DK1042 but *amyE*::P<sub>hyperspank</sub>-*mKate2* (Cm<sup>R</sup>) *sacA*::P<sub>tapA</sub>-*gfp* (Km<sup>R</sup>) | (50) |

TB852 was obtained by transforming DK1042 with gDNA isolated from TB601 and TB863, and selecting for tetracycline and kanamycin-resistant colonies, respectively. Strains TB514 and TB515 were obtained by transforming TB34 and TB35 respectively, with gDNA obtained from TB601 and TB602 and selecting for tetracycline and spectinomycin resistance. Cm<sup>R</sup>, Spec<sup>R</sup>, Km<sup>R</sup>, and Tet<sup>R</sup> denote chloramphenicol, spectinomycin, kanamycin, and tetracycline resistance cassettes, respectively.

### Structural heterogeneity assay

To assess structural heterogeneity of biofilms, pellicles were collected and transferred into a 1.5-mL microcentrifuge tube containing 1 mL of 0.9% (w/v) NaCl and ca. 20 µL of sterile glass sand ranging from 0.25-0.5 mm grain size (Carl Roth). Next, pellicles were vortexed (Scientific Industries, Vortex-Genie 2) for 2 min at 3200 rpm (maximal speed) and pellicle debris was allowed to sediment for 5 minutes. The dispersible fraction was transferred to a new Eppendorf tube, while the non-dispersible ‘clumps’ fraction was diluted in 1 mL of 0.9% (w/v) NaCl. Both fractions were sonicated as described previously (15), after which CFU levels were determined.

### Fitness assays

To determine the fitness costs of EPS and TasA production, mKATE2-labeled wild-type strains were competed with various biofilm-formation mutants. Overnight cultures were adjusted to the same optical density (OD), mixed at a 1:1 ratio, and 1% coculture inoculum was transferred into 1.5 mL MSgg medium. Cocultures were grown in static conditions at 30°C. CFU levels in both sonicated pellicle and liquid medium were determined immediately after inoculation and after 24 or 48 h of growth. Wild-type colonies were distinguished from biofilm mutants based on pink color (visible emission from mKate2 reporter). The selection rate (r) was calculated as the difference in the Malthusian parameters of both strains: r = ln[mutant (t=1)/mutant (t=0)]-(ln[wild-type(t=1)/wild-type(t=0)]), where t=1 is the time point at which the pellicle was harvested (51).

### Flow cytometry

Analysis was performed immediately after collection of the samples. To analyze expression levels of the *epsA-epsO* and *tapA-sipW-tasA* operons, flow cytometry analysis was performed using a BD FACScanto II (BD Biosciences). 100 000 cells per each sample were counted, where GFP+ were detected by blue laser (488) via 530/30 and mKate+ cells were detected by red laser (633) and 660/20 filter, respectively. Three replicates per condition were incubated at 30°C for 12, 16, 20, 24, 36, 40, 44, or 48 h. Afterwards, pellicles were harvested and sonicated. To study structural heterogeneity, harvested pellicles were vortexed as previously described, before sonication. Pellicles that were 12, 16, 20 or 24 h-old were diluted 20 times, whereas pellicles that were 36, 40, 44, or 48 hours old were diluted 200 times before flow cytometry analysis was performed. To obtain the average distribution of expression levels between replicates, data obtained from each replicate were subjected to binning using an identical bin size. Next, a mean count for each bin was obtained by averaging individual counts within this bin across all replicates, resulting in the mean distribution of single-cell level expression per condition.

### Microscopy and image analysis

To observe how biofilm mutants affect biofilm development, time lapse microscopy experiments were performed. Overnight cultures were adjusted to the same optical density (OD), mixed in a 1:3 ratio (wild-type:mutant), and inoculated in 500 µL MSgg medium inside an 8-well tissue culture chamber at 30°C (Sarstedt; width: 24 mm, length: 76 mm, growth area: 0.8 cm^2^). Bright-field images of pellicles were taken with an Axio Zoom V16 stereomicroscope (5x magnification; Carl Zeiss, Jena, Germany) equipped with a Zeiss CL 9000 LED light source, and an AxioCam MRm monochrome camera (Carl Zeiss), in which exposure time was set to 35 ms and images were captured every 15 minutes for a total of 48 h. Additionally, time-lapse videos of the wild-type monoculture biofilm development were recorded. For quantitative assessment of phenotypic heterogeneity, P_*eps*_*-gfp* pellicles were analyzed using a confocal laser scanning microscope (LSM 780, Carl Zeiss) equipped with a Plan-Apochromat/1.4 Oil DIC M27 63× objective and an argon laser (excitation at 488 nm for green fluorescence and 561 nm for red fluorescence, emission at 528 (±26) nm and 630 (±32) nm respectively). Zen 2012 Software (Carl Zeiss) and FIJI Image J Software (52) were used for image recording and subsequent processing, respectively.

Confocal microscopy images were used to extract the single-cell level distribution of *eps-gfp* expression using our recently developed BiofilmQ software (53). This analysis involved the registration of image time series to avoid sample drift, followed by top-hat filtering to eliminate noise, and Otsu thresholding to obtain a binary segmented image that separates the biofilm 3D location from the background. The BiofilmQ-inbuilt technique was used for dissecting this 3D volume into pseudo-cell cubes, which have the same volume as an average *B. subtilis* cell (4.6μm^3^) (54), based on the mKate fluorescence (constitutively expressed in all cells). Next, we quantified the GFP signal per pseudo-cell and plotted its distribution at different time points.

### Statistical analysis

Statistical differences between two experimental groups (e.g. total CFU/ml in robust biofilm fraction vs total CFU/ml in fragile biofilm fraction at single timepoint) were assessed using Two-Sample t-Test assuming equal variance. One-way ANOVA and Tukey Test was used for multiple samples comparisons (e.g. robust biofilm fraction across all sampling timepoints, where destructive sampling was applied). Two-way ANOVA and Tukey Test was used to assess the effects of time and different mutant on the WT in pellicle and in liquid fraction. Correlations were assessed using Pearson correlation coefficient. No statistical methods were used to predetermine sample size and the experiments were not randomized. All statistical tests were performed using OriginPro 2018 software.

## ACKNOWLEDGEMENTS

This study was supported by the Deutsche Forschungsgemeinschaft (DFG) to Á.T.K. (KO4741/2.1) within the Priority Program SPP1617, and the Collaborative Research Center SFB987 (to K.D.). S.B.O. and M. M. were supported by an Erasmus+ fellowship and a FEMS Research and Training Grant (FEMS-RG-2017-0054), respectively. This project has received funding from the European Union’s Horizon 2020 research and innovation programme under the Marie Skłodowska-Curie grant agreement No 713683 (H.C. Ørsted COFUND to A.D.) and the European Research Council (StG-716734 to K.D.). Work in the laboratory of Á.T.K. is partly supported by the Danish National Research Foundation (DNRF137) for the Center for Microbial Secondary Metabolites.

## AUTHOR CONTRIBUTIONS

Á.T.K. and A.D. conceived the project; S.B.O., M.M., D.S., R.H., and A.D. performed experiments; K.D. and S.B. contributed methodologies and equipment, respectively; S.B.O, A.D. and Á.T.K. wrote the manuscript, with all authors contributing to the final version.

