## Supplementary material for "Privatization of biofilm matrix in structurally heterogeneous biofilms": Fig S1 to 3

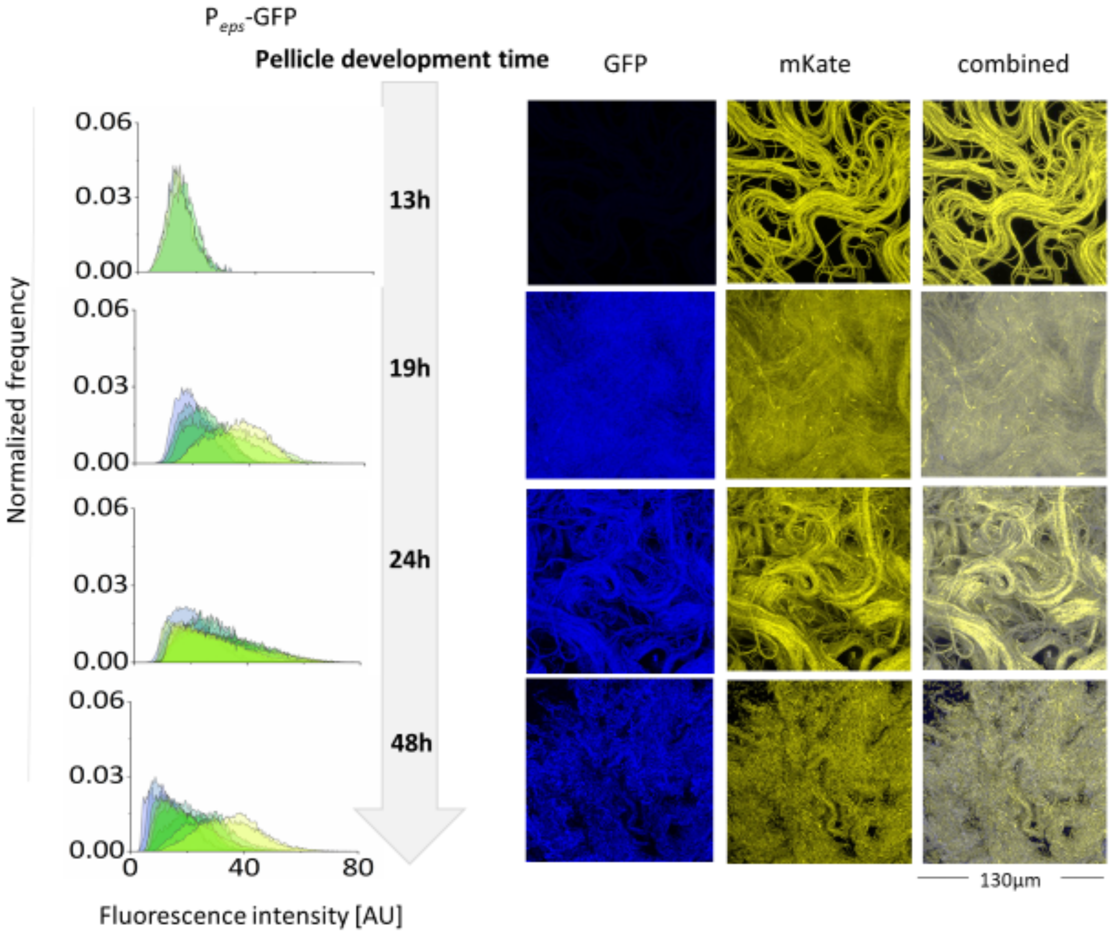


**FIG S1** Changes in matrix gene expression during biofilm development assessed by image analysis of intact pellicles. Signal intensity of $P_{eps}$-GFP was measured in each frame and represented as histogram. Histograms obtained from all frames (n=3 up to 9) are overlaid for each timepoint. On the right, representative confocal microscopy images: cells expressing $P_{epsA}$-GFP are colored blue, and cells constitutively expressing mKate2 are colored yellow. Z-stack with maximal pixel values for GFP channel was selected for representation.

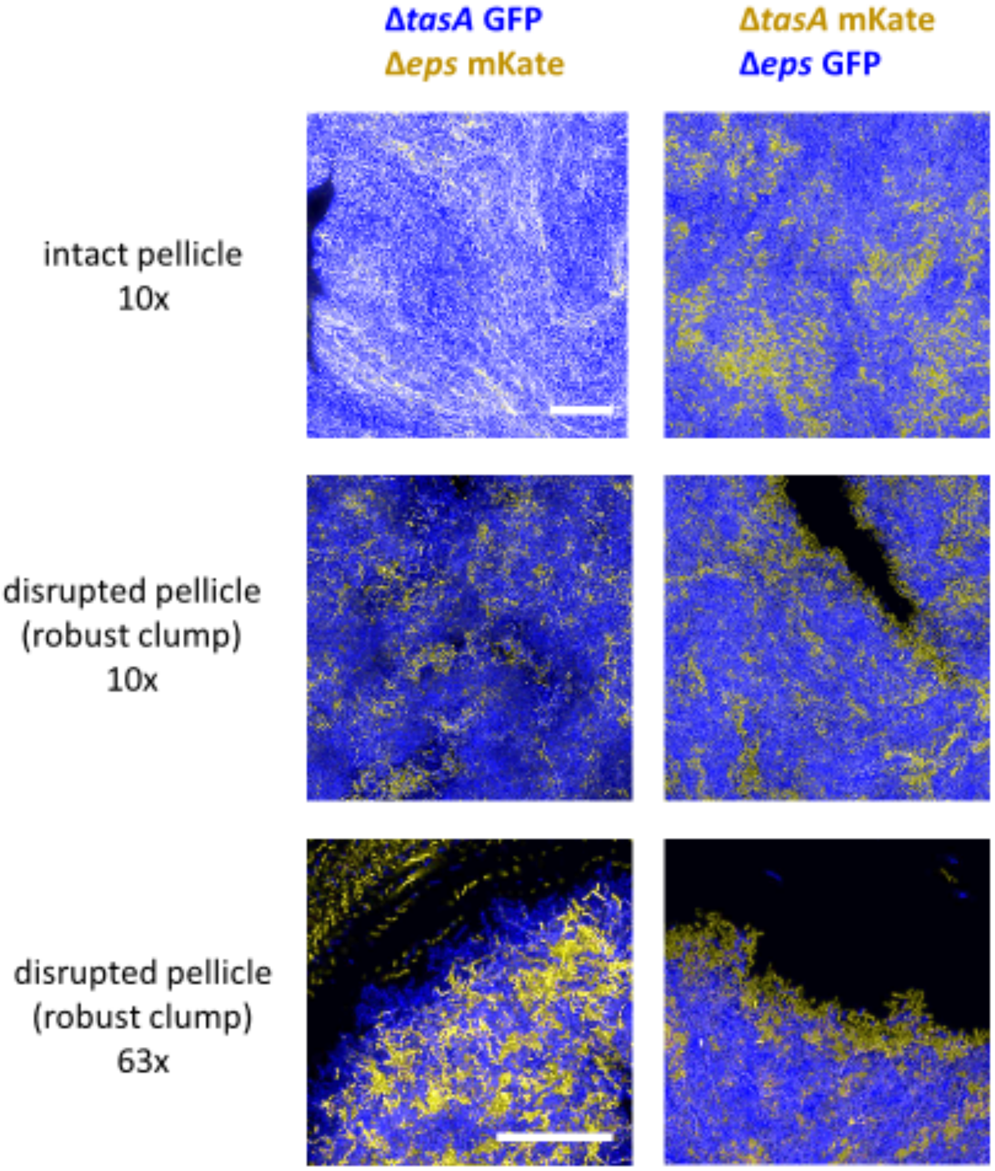


**FIG S2** Confocal images of mechanically disrupted and intact pellicles. Pellicles were captured at 48 hours using a confocal microscope. Upper panels represent intact pellicles formed by Δ*tasA* and Δ*eps* mutants at 10x magnification, middle panels represent identical mechanically disrupted pellicles and bottom panels represent identical mechanically disrupted pellicles at 63x magnification. Cells expressing mKate2 are colored in yellow while GFP producing cells are colored blue. Left and right images show opposing combinations of fluorescent markers. Scale bar: 50 µm.

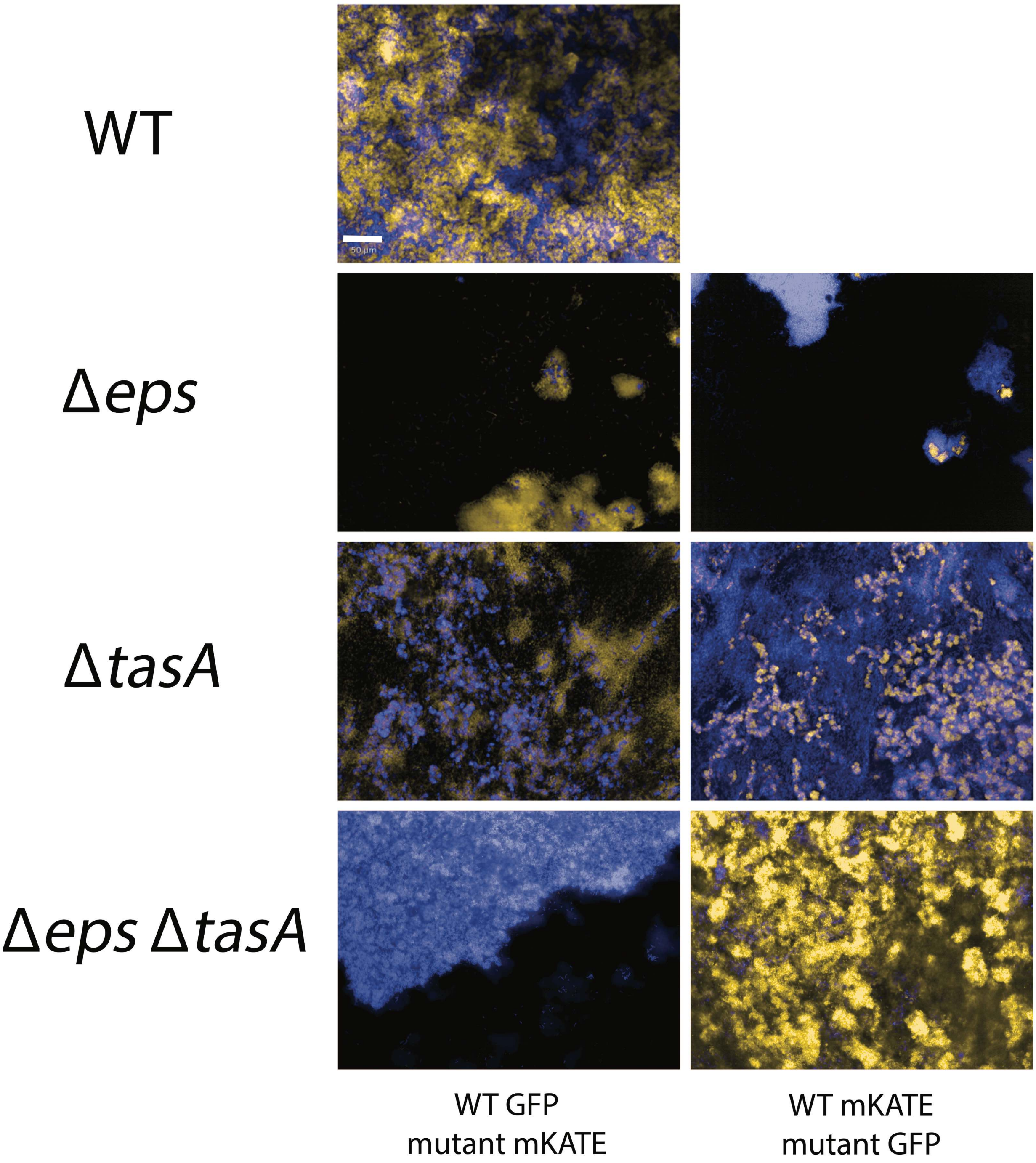


**FIG S3** Confocal microscopy images of mature pellicles (48h), initiated from co-culture of wild-type and different mutant strains, labelled with GFP (artificially coloured blue) and mKate (artificially coloured yellow). As a control, two wild-type strains labelled with different fluorescent markers were used. Scale bar: 50 µm.

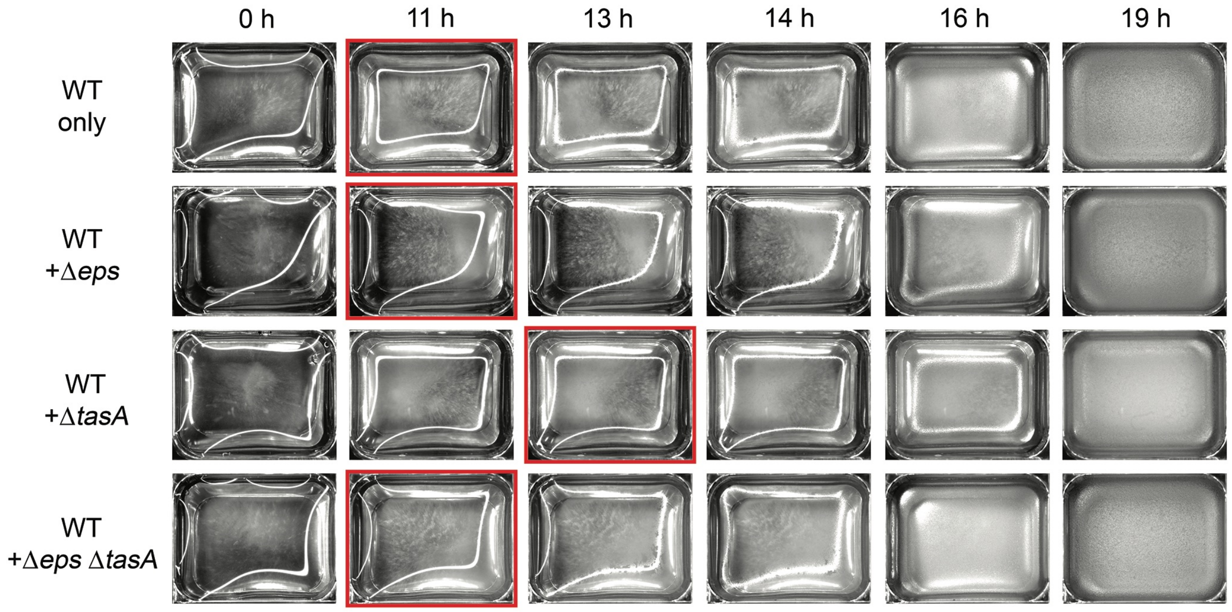


**FIG S4** Stereomicroscope images of pellicle formation. Pellicle formation images captured at various time points of WT, and co-cultures of WT + Δ*eps*, WT + Δ*tasA* or WT + Δ*eps*Δ*tasA* in MSgg medium incubated at 30°C inside an 8-well tissue culture chamber (Sarstedt; the wells have 24mm width and 76mm length, the growth area is 0.8 cm2). The red square represents the beginning of pellicle formation as indicated by disruption of the light reflection caused by the bright field stereomicroscope, which was assessed by close examination of the obtained images.

**Video S1. Stereomicroscope time lapse of pellicle formation.** Pellicle formation time lapse from images captured every 15 minutes for a total of 48 hours of identical WT, and WT + Δ*eps*, WT + Δ*tasA* or WT + Δ*eps*Δ*tasA* cocultures as described in Fig. S1. Cocultures were inoculated from top left to bottom right as follows: WT + Δ*eps*Δ*tasA*, WT only, WT + Δ*eps* and WT + Δ*tasA*. Frame rate: 10 frames per second. The wells have 24mm width and 76mm length.

**Supplemental dataset 1.** Raw data related to statistical analysis with corresponding p values.
